# Smallpox Vaccine Elicits CD8^+^ T Cell Immunity Persisting Over Four Decades with Cross-Recognition of Monkeypox Virus

**DOI:** 10.64898/2026.09.08.750003

**Authors:** Yujuan Du, Zongmian Yang, Yinying Xiao, Huajun Duan, Hong Kong, Chanchan Xiao, Jun Su

## Abstract

Monkeypox, caused by MPXV, has become a global public health concern; however, the T-cell immunity against MPXV following smallpox vaccination remains unclear. In this study, we screened the immunogenicity of 21 predicted MPXV epitopes and compared immune responses between young and elderly groups. Two immunogenic CD8⁺ T-cell epitopes, M15 and M17, were identified, both of which effectively stimulated specific immune responses, with significantly stronger responses observed in the young group. Notably, this study provides the first evidence that smallpox vaccination induces CD8⁺ T-cell immunity that cross-recognizes the MPXV epitopes M15 and M17 and remains detectable for more than four decades after vaccination, as demonstrated in vaccinated individuals aged 53–75. These findings provide experimental evidence for understanding the cross-protective mechanisms of the smallpox vaccine and informing monkeypox vaccine development.

## Introduction

Mpox is a zoonotic viral disease caused by Monkeypox virus (MPXV), belonging to the genus Orthopoxvirus in the family Poxviridae ^[1]^. MPXV was first identified in laboratory monkeys in 1958, with rodents likely serving as its natural reservoir ^[2]^. Human cases were first reported in Africa in the 1970s, followed by sustained transmission across the African continent ^[3]^. Since May 2022, a global mpox outbreak has affected multiple countries, prompting the World Health Organization to declare it a public health emergency of international concern. ^[4]^. Mpox has an incubation period of 5–21 days, with clinical manifestations including fever, headache, lymphadenopathy, and characteristic skin rash, and a case fatality rate of 0–11% ^[5, 6]^. Currently, no specific vaccine against mpox is available, and treatment primarily relies on supportive care; tecovirimat and other smallpox therapeutics are used only in specific outbreak control settings ^[7, 8]^.

The genus Orthopoxvirus comprises Variola virus (VARV), Vaccinia virus (VACV), and MPXV, all characterized by large linear double-stranded DNA genomes of approximately 190–197 kb ^[9]^. The central genomic region of MPXV shares 96.3% nucleotide homology with VARV, with amino acid sequence similarities of 91.7–99.2% for replication-associated proteins and 83.5–93.6% for terminal virulence proteins and immunomodulatory factors ^[10]^. This high degree of homology provides the molecular basis for cross-immunoprotection conferred by smallpox vaccination. Notably, 94% of CD4⁺ T cell epitopes and 82% of CD8⁺ T cell epitopes are shared between MPXV and VACV ^[11]^, suggesting that VACV-induced T cell immunity may elicit robust cross-reactivity against MPXV.

Anti-MPXV immune responses encompass both innate and adaptive immunity. Monocytes and neutrophils serve as initial targets of poxvirus infection, with early antigen detection predicting disease progression ^[12]^; natural killer cells also contribute to adaptive immune responses ^[13]^. Within adaptive immunity, B cells produce antibodies, CD4⁺ T cells provide helper functions, and CD8⁺ T cells play a critical role in viral clearance through cytotoxic killing of infected cells. Following smallpox vaccination, memory B cells and antibodies can persist for over 50 years; however, protective neutralizing antibody titers are maintained in only approximately 50% of vaccinees after 20 years ^[14]^. Memory CD4⁺ T cells exhibit an estimated half-life of 8– 15 years, with CD8⁺ T cells demonstrating comparable longevity ^[15]^. Importantly, SIV-infected macaques with CD4⁺ T cell counts below 300 cells/mm³ succumbed to MPXV challenge^[16]^, whereas HIV-infected individuals with counts exceeding 700 cells/mm³ experienced non-severe mpox ^[17]^, underscoring the antibody-independent role of T cells in modulating disease severity.

MHC tetramer technology is the gold standard method for quantitative detection of antigen-specific CD8⁺ T cells ^[18]^. This study employed this technique to screen MPXV-specific CD8⁺ T cell epitopes in order to evaluate cross-immunoprotection induced by smallpox vaccination. Human leukocyte antigens (HLA), the human MHC molecules, display extensive polymorphism across populations, with HLA-A2 being the most prevalent class I HLA allele in the Chinese population ^[19]^. Epitope presentation is MHC-restricted: endogenous antigens are proteasomally degraded, transported by TAP into the endoplasmic reticulum, and assembled with MHC-I molecules into pMHC complexes for surface presentation to CD8⁺ T cells ^[20]^. Bioinformatics tools such as IEDB enable prediction of MHC-I-restricted epitopes ^[21]^, while the T2A2 cell line, deficient in endogenous TAP expression, serves as an ideal platform for screening exogenous antigenic epitopes. Given the high homology between MPXV and VARV and the high frequency of HLA-A2 in the Chinese population, this study aims to identify MPXV HLA-A2-restricted CD8^+^ T cell epitopes and evaluate smallpox vaccine-induced cross-immunoprotection. Accordingly, the predicted peptides were subjected to comprehensive evaluation of pMHC binding, antigen presentation, tetramer recognition, T cell activation, cytotoxicity, and cross-reactive immunity elicited by smallpox vaccination, as illustrated in Figure 1.

**Fig. 1:**
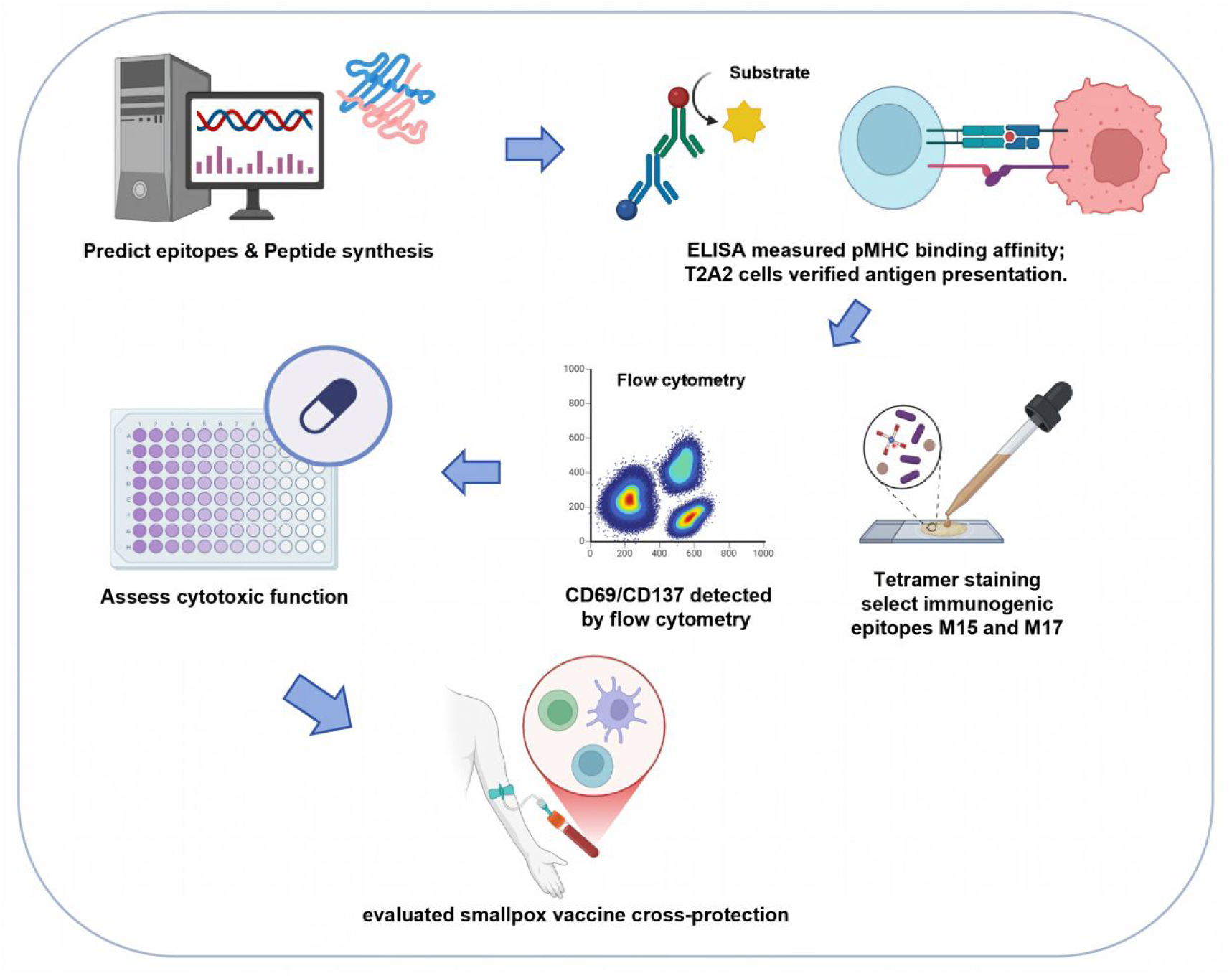
Workflow for screening and validating monkeypox virus CD8⁺ T-cell epitopes. Predicted peptides were evaluated for pMHC binding, antigen presentation, tetramer recognition, T-cell activation, cytotoxicity, and smallpox vaccine-induced cross-reactive immunity.

## Results

### Screening and Identification of Monkeypox Virus CD8⁺ T Cell Epitopes

First, bioinformatics prediction was performed using the IEDB database to analyze the 100% homologous sequences shared between monkeypox virus and smallpox virus (Figure 2 A). A large number of epitope predictions were obtained, from which the top twenty-one epitopes ranked by antigen presentation score were selected, with detailed information listed in Table 1.

**Fig. 2:**
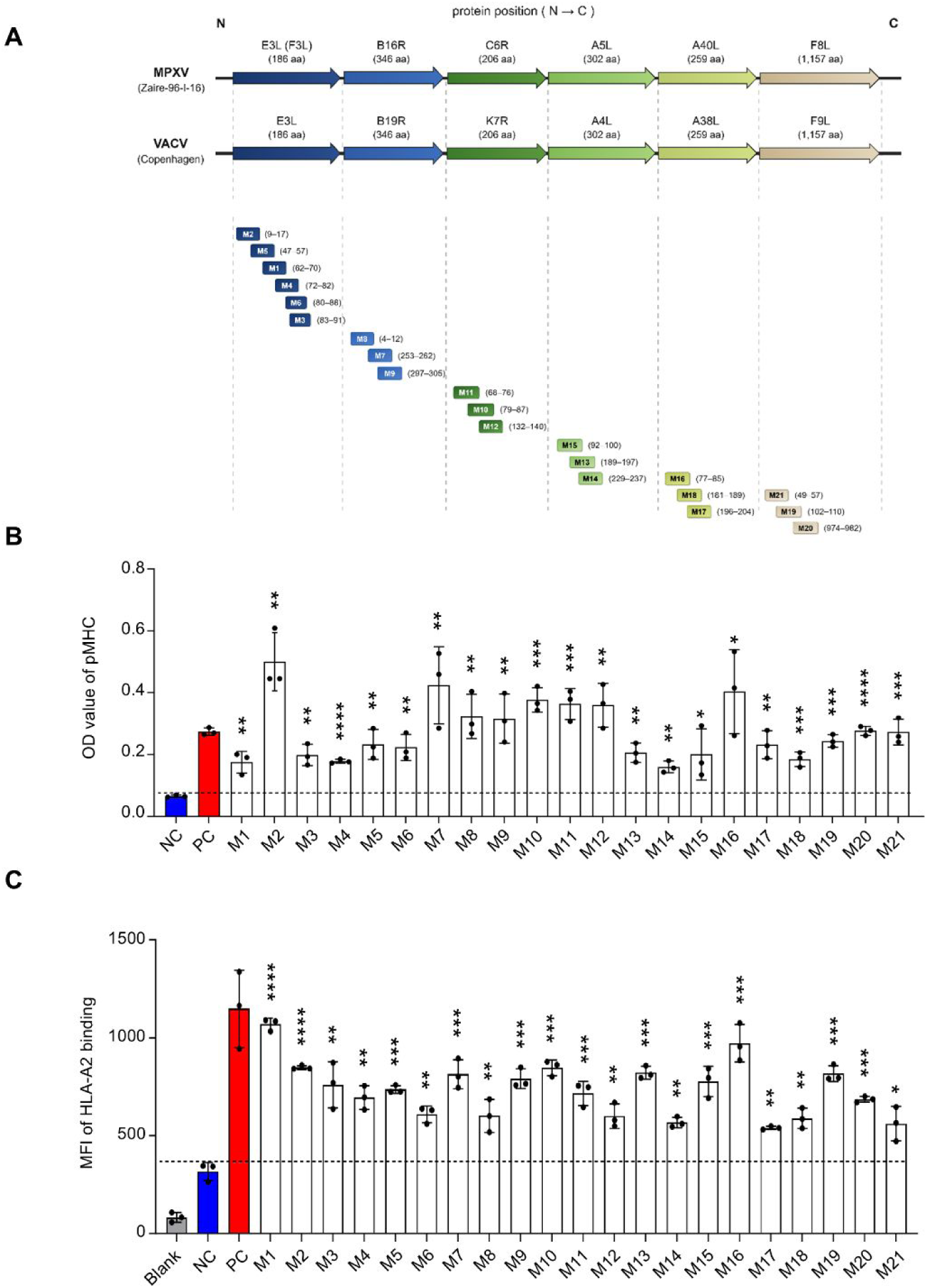
pMHC affinity determination and T2A2 cell stability analysis. **(A)** Schematic illustration of 21 candidate CD8^+^ T-cell epitopes (M1–M21) in six 100% homologous MPXV and VACV proteins. Epitopes are arranged from the N-terminus to the C-terminus of each protein, with amino-acid positions in the corresponding MPXV proteins shown in parentheses. **(B)** ELISA-based assessment of soluble pMHC complex formation and stability. Intact soluble HLA-A2/β2m/peptide complexes were detected using an HRP-conjugated anti-human β2m antibody, with absorbance at 414 nm as the readout. Data are shown as mean ± s.d., n = 3 independent experiments for each tested epitope. \*\*\*\**P*<0.0001, \*\*\**P*<0.001, \*\**P*<0.01, \**P*<0.05. two-sided t-test, comparing to NC. **(C)** T2A2 cell-based assessment of peptide-induced pMHC stabilization. Following peptide loading, cell-surface HLA-A2/β2m complexes were stained with a PE-conjugated anti-human β2m antibody and quantified by flow cytometry based on mean fluorescence intensity (MFI). Data are shown as mean ± s.d., n = 3 independent experiments for each tested epitope. \*\*\*\**P*<0.0001, \*\*\**P*<0.001, \*\**P*<0.01, \**P*<0.05. two-sided t-test, comparing to NC.

**Table 1.**
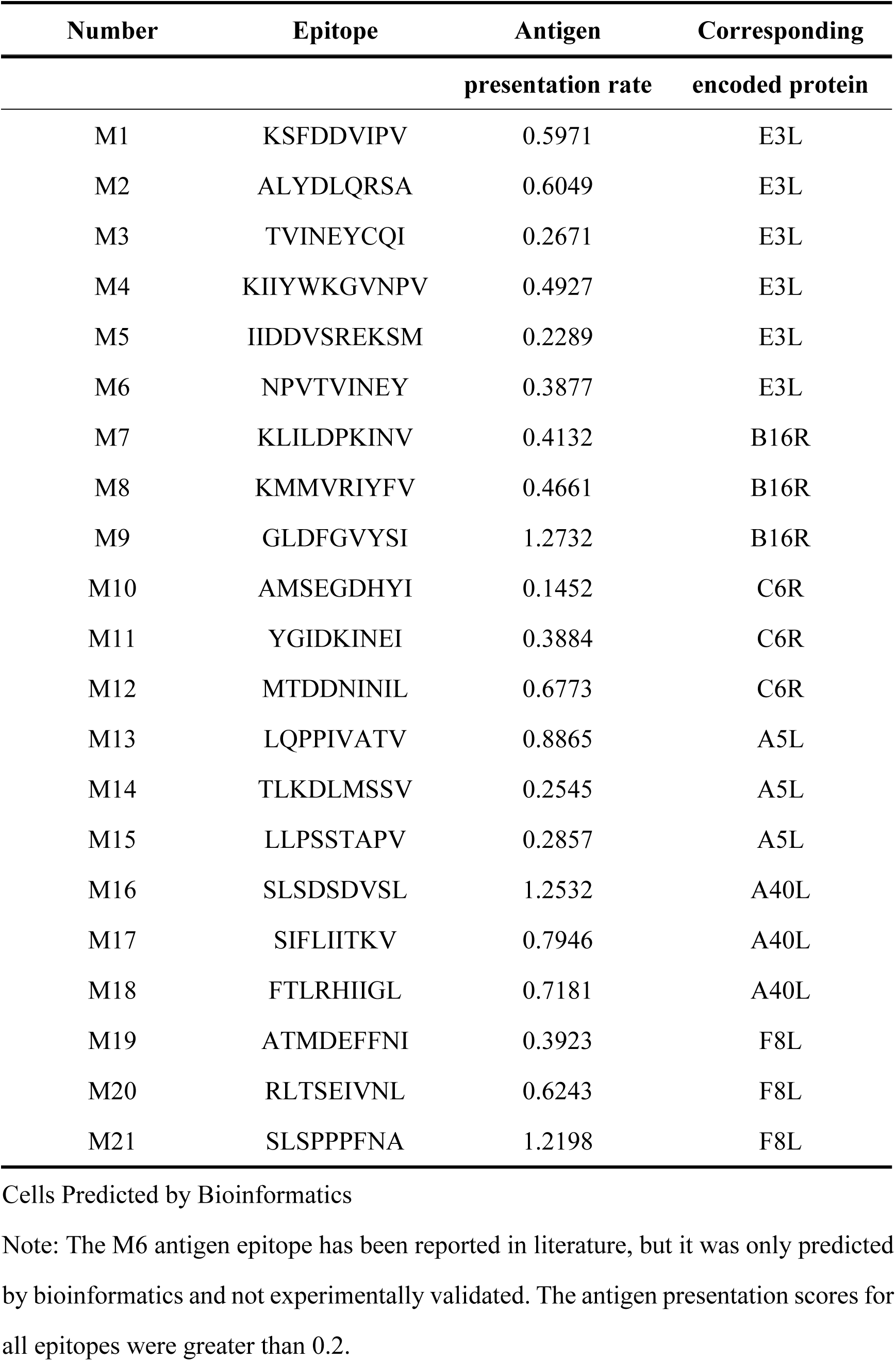
Antigenic Epitopes of Monkeypox Virus for HLA-A2-Restricted CD8^+^ T cells.

Subsequently, the corresponding antigenic peptides were custom-synthesized in collaboration with GenScript (Nanjing, China). A total of 21 HLA-A2-restricted CD8⁺ T cell epitopes (M1–M21) were identified, derived from viral-encoded proteins including E3L, B16R, C6R, A5L, A40L, and F8L. ELISA-based pMHC binding affinity assays demonstrated that all predicted epitopes could form stable complexes with HLA-A2 molecules, among which M2, M7, M8, M9, M10, M11, M12, and M16 exhibited binding affinities higher than that of the positive control peptide (Figure 2 B). Furthermore, antigen presentation capability was validated using the T2A2 cell line; flow cytometric analysis revealed that all 21 epitopes could form stable pMHC complexes on the cell surface with HLA-A2 (Figure 2 C), indicating effective presentation of these antigenic peptides. These results confirm that the predicted epitopes possess favorable HLA-A2 binding affinity and antigen presentation efficiency, laying a solid foundation for subsequent T cell activation experiments to screen for immunogenic epitopes.

### Activation and Detection of Monkeypox Virus Antigen-Specific CD8⁺ T Cells

The aforementioned screening only identified epitopes capable of successful antigen presentation; their immunogenicity remained to be validated through T cell activation experiments. We recruited six healthy volunteers and divided them into a young group (unvaccinated against smallpox) and an elderly group (born before 1980 and vaccinated against smallpox). Peripheral blood mononuclear cells (PBMCs) were isolated from venous blood, and CD8⁺ T cells were purified using magnetic beads. T2A2 cells, pre-treated with mitomycin and labeled with CFSE, were loaded with antigenic peptides and co-cultured with CD8⁺ T cells at a 1:1 ratio in 96-well plates in the presence of anti-CD28 antibody. After 16 hours, the expression of T cell activation markers CD69 and CD137 was detected.

CD69, an immediate early activation marker, showed significantly higher expression levels in the young group compared to the elderly group (*P* < 0.05) (Figure 3 A), indicating stronger CD8⁺ T cell activation responses in young individuals, which may be associated with age-related immune senescence. Additionally, since all elderly participants had received smallpox vaccination, this may also represent a contributing factor to their differential activation responses. CD137 (4-1BB), a critical co-stimulatory molecule playing key roles in T cell activation and proliferation, was similarly significantly higher in the young group than in the elderly group (*P* < 0.05) (Figure 3 B), further confirming the advantage of young individuals in immune responsiveness.

**Fig. 3:**
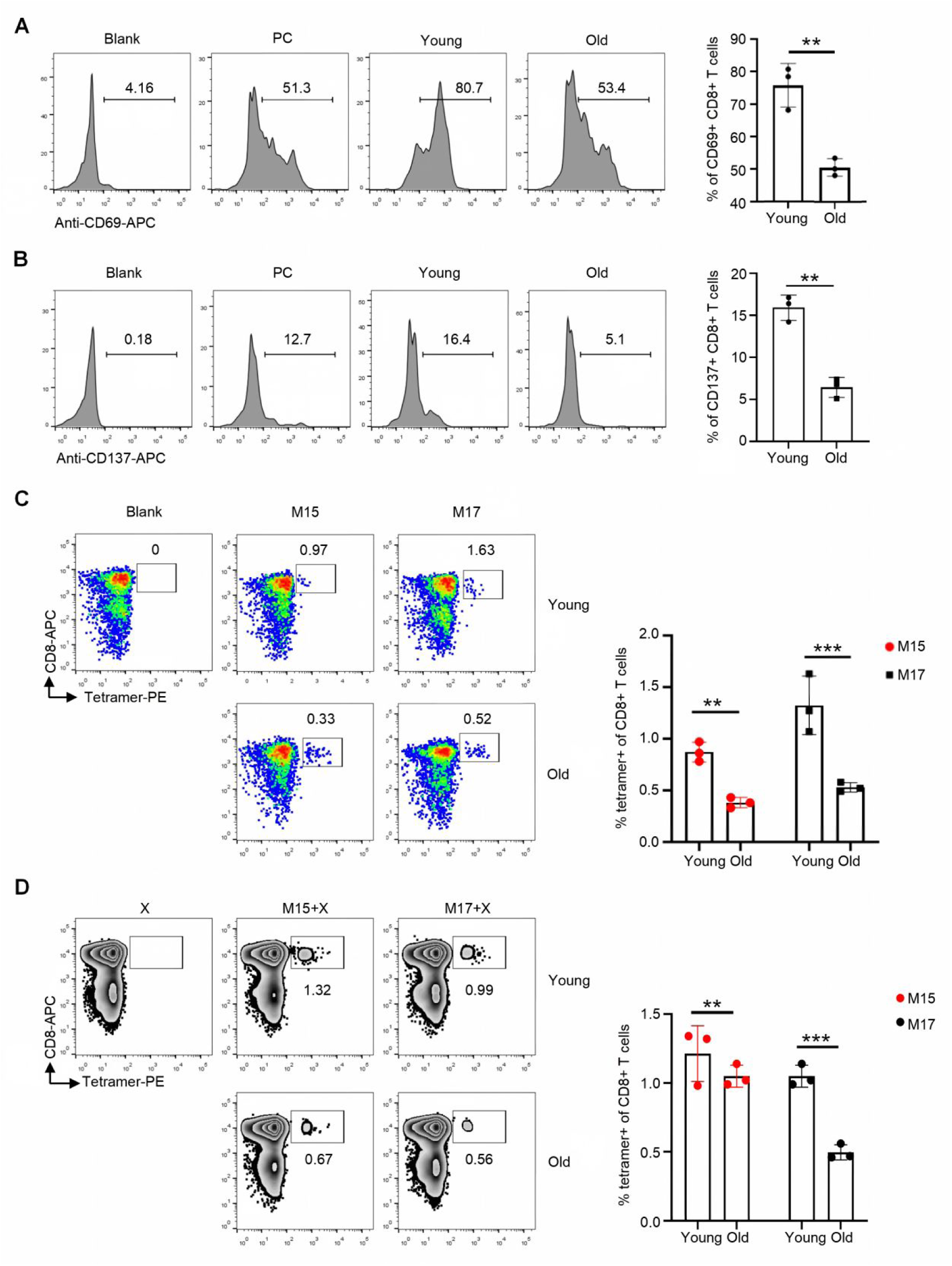
CD8^+^ T Cell Activation and Antigen-Specific Response Analysis. **(A)** Expression Level of CD69 in CD8^+^ T Cells. Data are shown as mean ± s.d., n = 3 independent experiments for each tested epitope. \*\**P*<0.01, two-sided t-test, the old group vs young group. **(B)** Expression Level of CD137 in CD8^+^ T Cells. Data are shown as mean ± s.d., n = 3 independent experiments for each tested epitope. \*\**P*<0.01, two-sided t-test, the old group vs young group. **(C)** After seven days of stimulation and activation, the tetramer staining results for antigen-specific CD8^+^ T cells showed positive peptides, along with corresponding statistical plots. Data are shown as mean ± s.d., n = 3 independent experiments for each tested epitope. *\*\*\*P*<0.001, \*\**P*<0.01. two-sided t-test, the old group vs young group. **(D)** Flow Cytometric Detection and Statistical Analysis of Tetramer Staining After Single Peptide Stimulation. Data are shown as mean ± s.d., n = 3 independent experiments for each tested epitope. *\*\*\*P*<0.001, \*\**P*<0.01. two-sided t-test, the old group vs young group.

To identify which antigenic epitopes specifically activated CD8⁺ T cells, tetramer staining was performed on activated CD8⁺ T cells on day 7 post-stimulation. Following stimulation, only M15 (LLPSSTAPV) and M17 (SIFLIITKV) showed positive staining (Figure 3 C), with the young group demonstrating significantly stronger responses than the elderly group (*P* < 0.05), consistent with the CD69 and CD137 findings. To validate single-peptide stimulation effects, we designed two peptide combinations: M15 combined with M1–M10, and M17 combined with M11–M20. Tetramer staining revealed that M15 induced higher numbers of antigen-specific CD8⁺ T cells than M17 (Figure 3 D), suggesting stronger immunogenicity of M15. Furthermore, the young group exhibited higher reactivity to both epitopes than the elderly group, demonstrating significant age-related differences, which may be attributed to the gradual decline of T cell function and immune responsiveness with aging.

### Cytotoxicity of Monkeypox Antigen-Specific CD8⁺ T Cells

To investigate the cytotoxic function of total CD8⁺ T cells in depth, we performed intracellular IFN-γ staining on activated total CD8⁺ T cells. IFN-γ is a key cytokine secreted primarily by activated total T cells and NK cells, playing an important role in antiviral immune responses. The results showed that IFN-γ expression levels in total CD8⁺ T cells were significantly higher in the young group than in the elderly group (Figure 4 A), suggesting more active T cell function in young individuals.

**Fig. 4:**
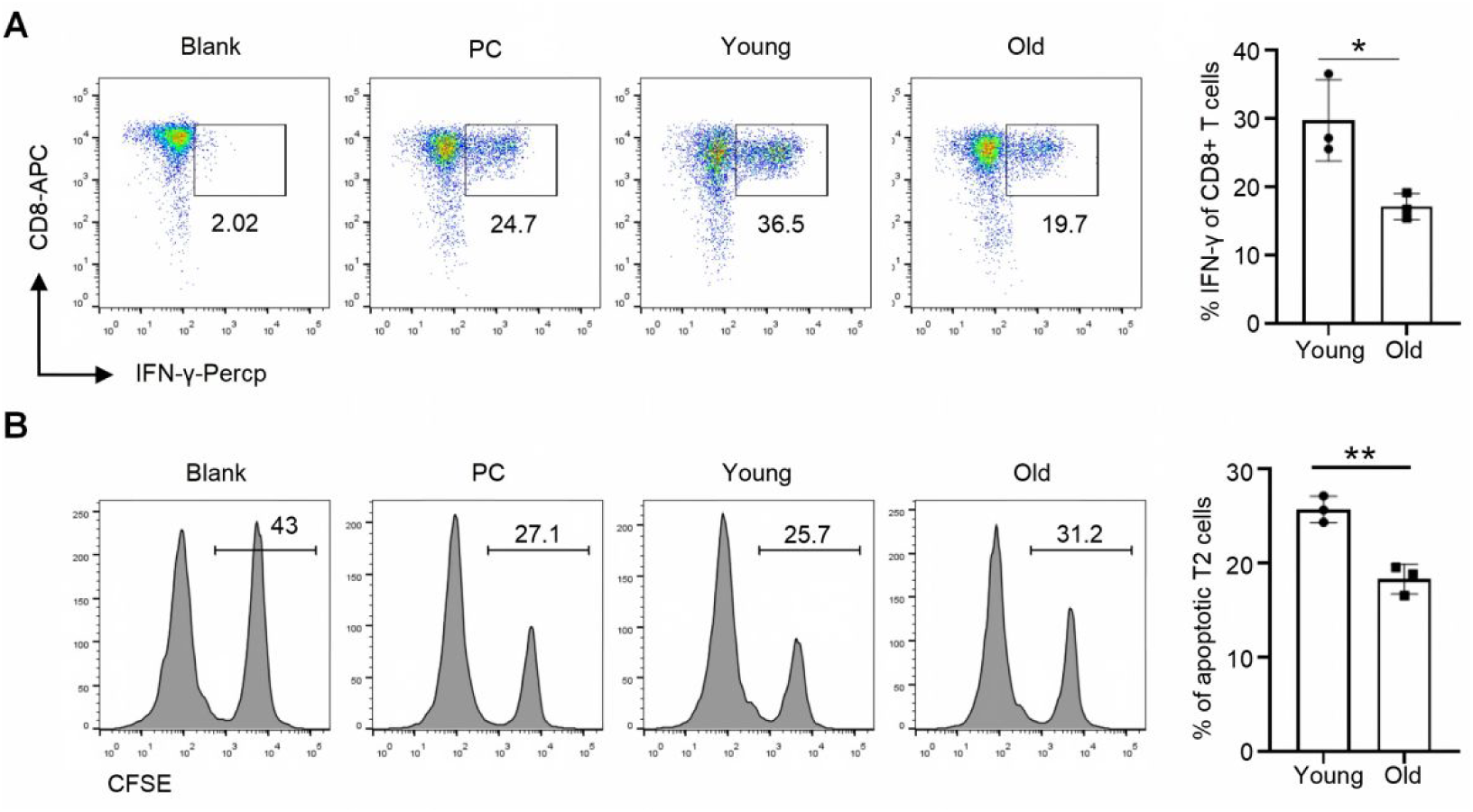
Functional Analysis of Total CD8^+^ T Cells. **(A)** Flow cytometric detection and statistical analysis of intracellular IFN-γ staining in total CD8^+^ T cells. Data are shown as mean ± s.d., n = 3 independent experiments for each tested epitope. \**P*<0.05, two-sided t-test, the old group vs young group. **(B)** Flow Cytometric Survival Assay and Killing Quantification of T2A2 Cells by CD8⁺ T Cells. Data are shown as mean ± s.d., n = 3 independent experiments for each tested epitope. \*\**P*<0.01, two-sided t-test, the old group vs young group.

Since T2A2 cells serve as both antigen-presenting cells and target cells for CD8⁺ T cells, we assessed their survival status by CFSE staining to reflect the killing efficiency of CD8⁺ T cells. The results demonstrated that the young group exhibited significantly higher killing efficiency against target cells than the elderly group (Figure 4 B), consistent with the intracellular IFN-γ findings. This age-related difference may be attributed to decreased T cell activation function, reduced secretion of cytotoxic cytokines, and diminished cytotoxicity in elderly individuals, indicating that older adults have relatively lower T cell immunity against monkeypox virus.

### Tetramer Staining for Detection of M15 and M17 Antigen-Specific CD8⁺ T Cells in Human Populations

To evaluate the cross-immunoprotective effect of smallpox vaccination against mpox, we utilized tetramer staining technology to detect CD8⁺ T cells specific for M15 and M17 antigens in different human populations. M15-and M17-specific HLA-A2 tetramers were prepared by UV-sensitive peptide exchange, and HLA-A2-positive individuals were analyzed by flow cytometry.

In unvaccinated individuals aged 0–46 years (n = 17), no M15-or M17-specific CD8⁺ T cells were detected (Figure 5 A).

**Fig. 5:**
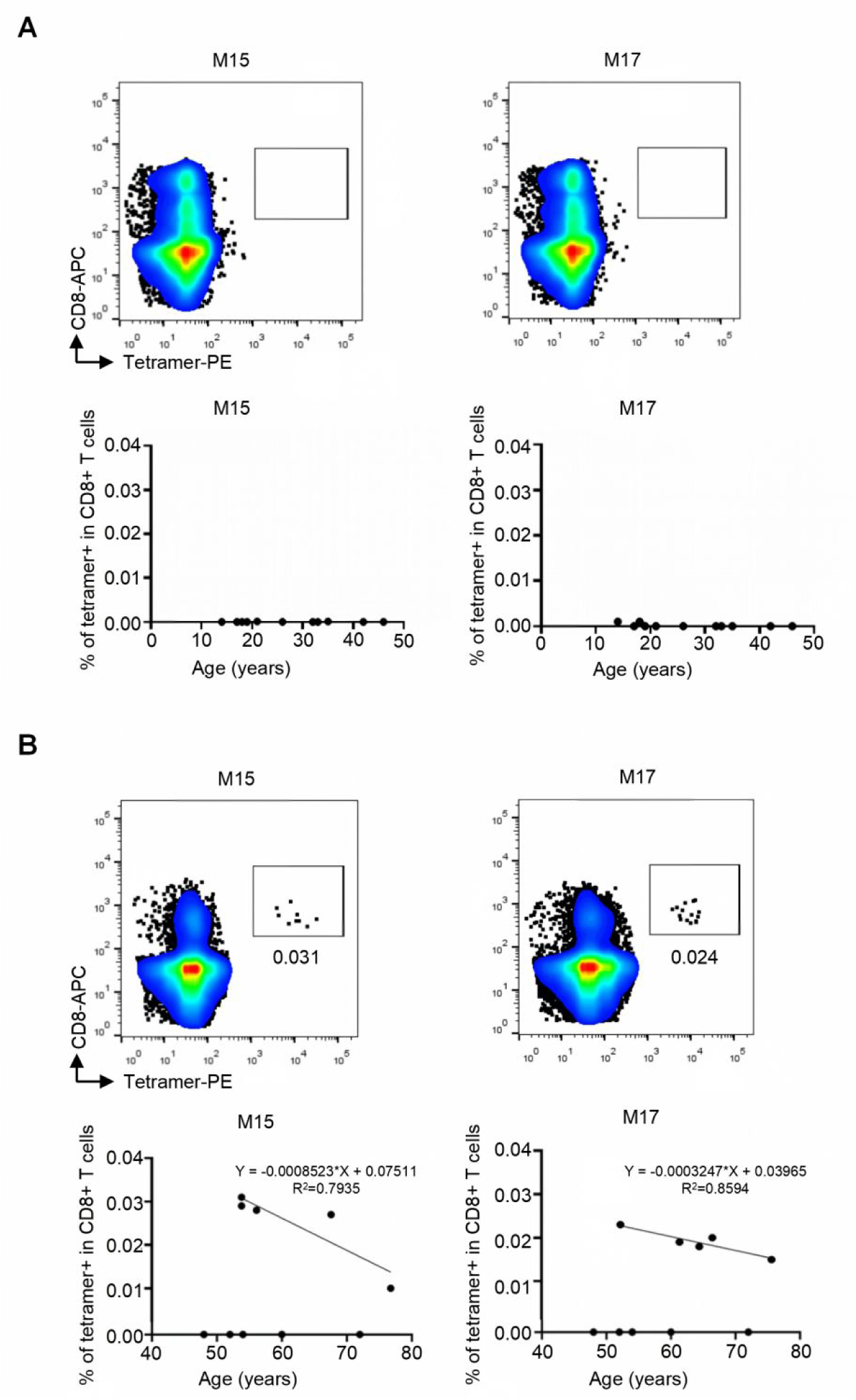
Tetramer Staining of Antigen-Specific CD8^+^ T Cells Against Smallpox. **(A)** Flow cytometry results of tetramer staining and statistical analysis in the unvaccinated population. **(B)** Flow cytometry results of tetramer staining and statistical analysis in the vaccinated population.

In smallpox-vaccinated individuals aged 48–75 years, cross-reactive CD8⁺ T cells were detected in 5 of 11 HLA-A2-positive individuals (45.5%), representing 18% (5/28) of the total vaccinated cohort subjected to tetramer analysis. with M15-specific CD8⁺ T cells accounting for 0.01%–0.031% and M17-specific cells for 0.015%–0.024% (Figure 5 B). Linear regression analysis revealed a significant age-dependent decline in cross-reactive T cell numbers (M15: Y = –0.0007321X + 0.06878, R² = 0.7851, F = 10.96, *P* = 0.0454; M17: Y = –0.0003687X + 0.04272, R² = 0.8856, F = 23.23, *P* = 0.017), indicating a significant negative correlation between age and the number of M15 and M17 epitope cross-reactive T cells.

## Discussion

The 2022 global MPXV outbreak was declared a public health emergency by WHO ^[22]^. MPXV, a zoonotic orthopoxvirus, spreads through contact with infected skin, fluids, or contaminated objects ^[23]^. Although monkeypox resembles smallpox clinically due to high genomic homology ^[24]^, its fatality rate is lower. Notably, reinfection is rare after clearance, likely due to functional immunological memory ^[25]^.

While human immune data remain limited, animal studies show CD8^+^ T cells play a predominant role in viral clearance ^[26]^. Smallpox-vaccinated individuals over age 45 possess MPXV-reactive CD8^+^ T cells, indicating cross-reactive immunity ^[27]^. However, as smallpox vaccination ended in the 1970s without boosters, cross-protection remains controversial and largely limited to older populations ^[28]^. No MPXV-specific vaccines exist, necessitating urgent vaccine development.

This study identified MPXV-specific CD8^+^ T cell epitopes and evaluated smallpox vaccine cross-protection. We selected MPXV-VACV homologous sequences and used IEDB to predict HLA-A2-restricted epitopes. MHC tetramer technology, a key method for identifying virus-specific CD8^+^ T cells ^[29]^, was employed for screening. Given that HLA-A2 is the most prevalent allele in Chinese populations, we prepared HLA-A2 tetramers using prokaryotic expression, nickel affinity purification, and refolding with photosensitive peptides ^[30]^. Size-exclusion chromatography yielded >95% pure monomers; UV-mediated peptide exchange and streptavidin assembly produced functional tetramers.

Grifoni et al. reported 94% of CD4 and 82% of CD8 epitopes are conserved between orthopoxviruses and MPXV, suggesting strong VACV-MPXV cross-reactivity. Rcheulishvili et al. designed a universal mRNA vaccine using conserved MPXV-VACV-VARV sequences, demonstrating MHC binding and immune induction ^[31]^. Most MPXV vaccine research remains computational; our experimental validation using a laboratory-developed screening system provides more robust evidence. Our 21 predicted epitopes, all from MPXV-VACV homologous regions, avoid poxvirus immune evasion interference and biosafety concerns.

All predicted epitopes showed significantly higher HLA-A2 binding affinity than controls, forming stable pMHC complexes. Using our artificial antigen presentation system with T2A2 cells, we identified M15 and M17 as immunogenic epitopes. Younger individuals showed stronger CD8^+^ T cell activation, IFN-γ production, and cytotoxicity than older individuals, consistent with the reported 8–15 years half-life of antiviral T cells. To our knowledge, this study provides the first evidence that smallpox vaccine–elicited CD8⁺ T-cell memory capable of cross-recognizing the MPXV epitopes M15 and M17 remains detectable for more than four decades after vaccination. These cross-reactive T cells were detected in vaccinated individuals aged 53–75, although their frequencies declined with age, indicating the remarkable durability of vaccine-induced cellular immune memory despite its gradual waning over time.

We only examined HLA-A2, the most common Chinese subtype, without assessing other alleles. The specific protective efficiency of smallpox vaccine against monkeypox was not quantified. Future studies should include additional HLA alleles and comprehensive cross-protection assessments.

## Materials and Methods

### Population Study Design

A total of 100 healthy volunteers were recruited, spanning ages 0–79 years. Subjects were divided into two groups: unvaccinated individuals aged 0–45 years (born after 1980) and vaccinated individuals aged 46–79 years (born before 1980). HLA-A2-positive individuals were identified by flow cytometry, and tetramer staining was performed on their PBMCs to detect M15-and M17-specific CD8⁺ T cells. Detailed volunteer information and HLA-A2 subtype identification are provided in Table S1. The institutional review board of Jinan University School of Medicine approved the study (JNUKY-2023-0049).

### Isolation of PBMCs

Healthy human peripheral blood was collected in heparinized vacutainers and kept at room temperature until processing. PBMCs were isolated by density gradient centrifugation using lymphocyte separation medium (Coolaber, China). Cell viability was assessed by Trypan blue staining. PBMCs were cryopreserved in fetal bovine serum with 10% DMSO (Sigma‒Aldrich, US) and stored in liquid nitrogen until use.

### HLA-A2 Typing

PBMCs were stained with PE-conjugated anti-human HLA-A2 antibody (BioLegend, US) after blocking with Human TruStain FcX (BioLegend, US) for 10 min at room temperature. Cells were incubated with the antibody for 20 min on ice, washed, and resuspended in cell staining buffer. 7-AAD viability staining solution (BioLegend, US) was added to exclude dead cells. Samples were analyzed by flow cytometry (BD FACSCanto II, US).

### Isolation of CD8⁺ T Cells

CD8⁺ T cells were negatively selected from PBMCs using the EasySep™ Human CD8⁺ T Cell Isolation Kit (STEMCELL, Canada) according to the manufacturer’s instructions. Briefly, cells were resuspended in EasySep buffer at a density of 5×10⁷ cells/mL, incubated with the isolation cocktail and RapidSpheres for 5 min at room temperature, and then subjected to magnetic separation. The enriched CD8⁺ T cells were collected and resuspended in T cell culture medium for subsequent experiments.

### Antigen Peptide Prediction and Synthesis

HLA-A2-restricted CD8⁺ T cell epitopes of monkeypox virus were predicted using the IEDB MHC-I prediction tool (http://tools.iedb.org/mhci/) based on sequence homology between monkeypox virus and vaccinia virus. The top 20 predicted epitopes with IC₅₀ < 100 nM were selected and synthesized (GenScript, China). A previously reported epitope (M6) and influenza peptide GILGFVFTL were used as controls ^[32]^.

### ELISA for pMHC Stability

Streptavidin-coated ELISA plates (BioLegend, US) were incubated with biotinylated HLA-A2 monomers containing test peptides. HRP-conjugated anti-human β2-microglobulin antibody (BioLegend, US) was added to detect intact pMHC complexes. After washing, the ABTS substrate solution was added, and absorbance at 414 nm was measured using a microplate reader (BioTek, US).

### T2A2 Cell Peptide Loading and Stability Assay

T2A2 cells were cultured in IMDM medium supplemented with 10% fetal bovine serum and antibiotics. For peptide loading, cells were incubated with 20 μM synthetic peptides for 4 h at 37 ℃. Cells were then stained with FITC-conjugated anti-human HLA-A2 antibody (BioLegend, US) and analyzed by flow cytometry. Mean fluorescence intensity (MFI) was used to evaluate pMHC complex stability.

### In Vitro Stimulation of CD8⁺ T Cells

T2A2 cells were pretreated with mitomycin C (20 μg/mL, GLPBIO, US) for 30 min and labeled with CFSE (TargetMol, US). Peptide-loaded T2A2 cells were co-cultured with autologous CD8⁺ T cells at a 1:1 ratio (5×10⁵ cells each per well) in 96-well plates in the presence of anti-human CD28 antibody (1 μg/mL, BioLegend, US) and IL-2 (50 IU/mL, Sigma‒Aldrich, US). Cells were restimulated with fresh peptide and IL-2 every two days. Activation markers CD69 and CD137 were detected at 16 h, and tetramer staining was performed on day 7.

### Tetramer Staining

HLA-A2 tetramers loaded with M15 (LLPSSTAPV) or M17 (SIFLIITKV) peptides were prepared. For staining, PBMCs or stimulated CD8⁺ T cells were incubated with Human TruStain FcX for 10 min at room temperature, followed by tetramer incubation for 1 h at 25°C in the dark. APC-conjugated anti-human CD8 antibody (BioLegend, US) was then added. After washing, 7-AAD was added to exclude dead cells. Samples were analyzed by flow cytometry within 2 h.

### Intracellular Cytokine Staining

On day 7 of stimulation, total CD8^+^ T cells were restimulated with leukocyte activation cocktail (BD Biosciences, US) containing PMA, ionomycin, and Brefeldin A for 6 h at 37℃. Cells were surface-stained with APC-conjugated anti-human CD8 antibody, fixed, and permeabilized. Intracellular IFN-γ was stained with PerCP-conjugated anti-human IFN-γ antibody (BioLegend, US) and analyzed by flow cytometry.

### Cytotoxicity Assay

The cytotoxic activity of antigen-specific CD8⁺ T cells was assessed by CFSE-labeled T2A2 target cell killing. After 7 days of co-culture, the percentage of surviving CFSE⁺ T2A2 cells was determined by flow cytometry. Cell death was quantified by 7-AAD uptake.

### Funding

This work was supported by the National Key R&D Program of China (2023YFE0118700); National Natural Science Cross disciplinary Major Research Program (92374203); the National Natural Science Foundation of China (82603446); Guangdong Medical Research Foundation Project (B2026256); the Fellowship of China Postdoctoral Science Foundation (2026T190386; 2025M781367); the Guangdong Basic and Applied Basic Research Foundation (2025A1515010448); Natural Science Foundation of Hunan Province of China(2026JJ60274); National Innovation and Entrepreneurship Training Program For Undergraduate (202610559047).

## Consent to participate

Informed consent was obtained from all participants or their legal guardians prior to enrollment.

## Data Availability

The data that support the findings of this study are available from the corresponding author upon reasonable request.

## Author contributions

Chanchan Xiao and Jun Su were responsible for the conception and overall design of the study, whereas Yujuan Du, Zongmian Yang and Yinying Xiao collaborated on conceptualization, data management, and methodology and contributed to the writing. In addition, Huajun Duan and Hong Kong performed the formal analysis and participated in the review and editing process of the manuscript.

## Supplementary Tables

**Table S1.** Volunteer information and identification of HLA-A2 subtypes.

| **Volunteer ID** | **Age** | **Gender** | **Routine blood test** | **HLA-A2 Positive** |
| --- | --- | --- | --- | --- |
| 1 | 11 months old | Woman | Normal | - |
| 2 | 12 years old | Man | Normal | - |
| 3 | 14 years old | Woman | Normal | - |
| 4 | 14 years old | Man | Normal | + |
| 5 | 16 years old | Man | Normal | - |
| 6 | 16 years old | Woman | Normal | - |
| 7 | 16 years old | Man | Normal | - |
| 8 | 16 years old | Man | Normal | - |
| 9 | 17 years old | Man | Normal | + |
| 10 | 17 years old | Woman | Normal | + |
| 11 | 17 years old | Woman | Normal | + |
| 12 | 18 years old | Man | Normal | - |
| 13 | 18 years old | Woman | Normal | + |
| 14 | 19 years old | Woman | Normal | + |
| 15 | 21 years old | Woman | Normal | - |
| 16 | 21 years old | Man | Normal | - |
| 17 | 21 years old | Man | Normal | + |
| 18 | 21 years old | Woman | Normal | + |
| 19 | 21 years old | Man | Normal | - |
| 20 | 22 years old | Woman | Normal | - |
| 21 | 22 years old | Man | Normal | - |
| 22 | 22 years old | Man | Normal | - |
| 23 | 22 years old | Woman | Normal | - |
| 24 | 22 years old | Woman | Normal | - |
| 25 | 26 years old | Woman | Normal | - |
| 26 | 26 years old | Woman | Normal | + |
| 27 | 26 years old | Man | Normal | - |
| 28 | 26 years old | Man | Normal | + |
| 29 | 26 years old | Man | Normal | - |
| 30 | 26 years old | Woman | Normal | - |
| 31 | 26 years old | Man | Normal | + |
| 32 | 26 years old | Man | Normal | - |
| 33 | 32 years old | Woman | Normal | - |
| 34 | 32 years old | Woman | Normal | - |
| 35 | 32 years old | Woman | Normal | + |
| 36 | 32 years old | Man | Normal | - |
| 37 | 32 years old | Woman | Normal | + |
| 38 | 33 years old | Man | Normal | - |
| 39 | 33 years old | Woman | Normal | - |
| 40 | 33 years old | Man | Normal | + |
| 41 | 35 years old | Woman | Normal | + |
| 42 | 35 years old | Man | Normal | - |
| 43 | 35 years old | Woman | Normal | - |
| 44 | 35 years old | Woman | Normal | - |
| 45 | 35 years old | Woman | Normal | - |
| 46 | 40 years old | Man | Normal | - |
| 47 | 40 years old | Woman | Normal | - |
| 48 | 42 years old | Man | Normal | + |
| 49 | 42 years old | Man | Normal | - |
| 50 | 43 years old | Man | Normal | - |
| 51 | 46 years old | Woman | Normal | - |
| 52 | 46 years old | Woman | Normal | + |
| 53 | 46 years old | Woman | Normal | - |
| 54 | 46 years old | Man | Normal | - |
| 55 | 46 years old | Man | Normal | - |
| 56 | 48 years old | Woman | Normal | - |
| 57 | 48 years old | Woman | Normal | - |
| 58 | 48 years old | Woman | Normal | + |
| 59 | 48 years old | Woman | Normal | + |
| 60 | 48 years old | Man | Normal | - |
| 61 | 51 years old | Man | Normal | - |
| 62 | 52 years old | Woman | Normal | - |
| 63 | 52 years old | Woman | Normal | + |
| 64 | 52 years old | Man | Normal | + |
| 65 | 54 years old | Woman | Normal | - |
| 66 | 54 years old | Woman | Normal | - |
| 67 | 54 years old | Woman | Normal | - |
| 68 | 54 years old | Man | Normal | - |
| 69 | 54 years old | Man | Normal | - |
| 70 | 54 years old | Woman | Normal | + |
| 71 | 56 years old | Woman | Normal | - |
| 72 | 56 years old | Woman | Normal | - |
| 73 | 56 years old | Man | Normal | - |
| 74 | 57 years old | Woman | Normal | - |
| 75 | 60 years old | Man | Normal | - |
| 76 | 60 years old | Woman | Normal | - |
| 77 | 60 years old | Man | Normal | + |
| 78 | 61 years old | Man | Normal | - |
| 79 | 61 years old | Woman | Normal | - |
| 80 | 61 years old | Woman | Normal | - |
| 81 | 61 years old | Woman | Normal | + |
| 82 | 61 years old | Woman | Normal | - |
| 83 | 62 years old | Woman | Normal | - |
| 84 | 62 years old | Man | Normal | - |
| 85 | 64 years old | Woman | Normal | + |
| 86 | 65 years old | Woman | Normal | - |
| 87 | 65 years old | Woman | Normal | - |
| 88 | 66 years old | Woman | Normal | - |
| 89 | 66 years old | Woman | Normal | + |
| 90 | 66 years old | Woman | Normal | - |
| 91 | 70 years old | Man | Normal | - |
| 92 | 70 years old | Man | Normal | - |
| 93 | 71 years old | Man | Normal | - |
| 94 | 71 years old | Woman | Normal | - |
| 95 | 72 years old | Woman | Normal | - |
| 96 | 72 years old | Woman | Normal | + |
| 97 | 75 years old | Woman | Normal | + |
| 98 | 75 years old | Woman | Normal | - |
| 99 | 77 years old | Man | Normal | - |
| 100 | 79 years old | Woman | Normal | - |

Note:
(1) The complete blood count parameters include: white blood cell count ( $3.5\text{--}9.5 \times 10^9/\text{L}$ ); red blood cell count ( $3.80\text{--}5.10 \times 10^9/\text{L}$ ); neutrophil percentage (0.4–0.75); lymphocyte percentage (0.2–0.5); monocyte percentage (0.03–0.1); eosinophil percentage (0.04–0.08); basophil percentage (0–0.06); neutrophil count ( $1.8\text{--}6.3 \times 10^9/\text{L}$ ); lymphocyte count ( $1.1\text{--}3.2 \times 10^9/\text{L}$ ); monocyte count ( $0.1\text{--}0.6 \times 10^9/\text{L}$ ); eosinophil count ( $0.02\text{--}0.52 \times 10^9/\text{L}$ ); basophil count ( $0\text{--}0.06 \times 10^9/\text{L}$ ).
(2) + indicates HLA-A2 positivity; – indicates HLA-A2 negativity.

